# DomDiff: protein family and domain annotation via diffusion model and ESM2 embedding

**DOI:** 10.1101/2025.10.28.685005

**Authors:** Chao Zhang, Haopeng Xia, Peng Yin

## Abstract

Accurate identification of conserved protein domain boundaries and their classification are fundamental to genome annotation, but are hindered by ambiguous boundaries, cross-domain interference, and limited samples for rare families. Here, we present DomDiff, a supervised conditional diffusion framework that reformulates the task as a generative process. Taking ESM2 embeddings, secondary structures, and biLSTM priors as inputs, it generates labels from Gaussian noise through iterative denoising, allowing coarse-to-fine optimization. We conducted a series of benchmark analyzes on publicly available protein sequence datasets, showing that DomDiff outperforms existing methods in domain boundary identification and classification, delivering performance gains of 12.6% in boundary detection and 4.2% in classification accuracy compared to other leading models. It excels particularly in annotating rare families, offering a powerful tool for specific applications such as large-scale genome annotation and functional characterization of novel proteins, thus providing a new paradigm for few-shot challenges in bioinformatics.

## 1 Introduction

As primary molecular constituents, protein functions mediate most intracellular biological processes, and various function domains are encoded within amino acid sequences[1]. The conserved domains are recognized as fundamental units of function, structure, and evolution[2]. Statistically, most amino acid sequences, particularly those of eukaryotes, are structured as concatenations of one or more conserved domains[3]. Consequently, the precise identification of conserved domain boundaries and functional classification of amino acid sequences constitute the cornerstone of genome annotation, protein function prediction, and evolutionary analysis.

However, accurately identifying conserved domain boundaries from amino acid sequences remains a formidable challenge, stemming primarily from three key factors. First, identifying the amino acid sequence of conserved domains is more complex than initially apparent. Conservation patterns diverge substantially across different protein families, and their boundaries are often fuzzy and dependent on global sequences[4][5][6]. Second, interference between distinct conserved domains significantly increases the difficulty of boundary identification[7]. Finally, the classification of rare domains constitutes a major bottleneck, despite the explosive growth of sequence data, many protein families are small or rare, resulting in limited samples for modeling[8].

Prior efforts to tackle this task have focused on amino acid sequence similarity, roughly falling into three categories: (i) methods based on multiple sequence alignment (MSA)[9][10] (ii) methods based on genealogical clustering[11][12][13][14] and (iii) methods based on a deep learning model[15][16][17][18][19][20]. For example, HMMER[10] builds MSA-based hidden Markov models (HMMs) for protein homology search and family annotation, but its heavy reliance on homologous sequences limits applicability to rare proteins. ProtoNet[21], a genealogical clustering method, organizes protein families through large-scale sequence similarity clustering. DeepFam[20] is a supervised convolutional neural network (CNN) with inherent data dependency; its performance depends on data scale and diversity, leading to overfitting for small-sample families and failure to capture generalizable family-specific features. DeepDom[17] uses bidirectional long-short-term memory (biLSTM) to model sequence contextual dependencies for domain boundary prediction, but inadequate long-range sequence modeling limits its boundary accuracy, particularly struggling with ambiguous boundaries in multidomain proteins due to cross-domain interference. These collective limitations highlight a clear gap for a new paradigm capable of holistically addressing boundary ambiguity, domain interference, and the long-tail challenge of rare families.

To address these challenges, we propose DomDiff, a novel supervised conditional diffusion-based framework that takes advantage of ESM2[22] embeddings to generate domain boundaries and categories of protein families from the noise sequence. DomDiff offers three key advantages as follows. First, DomDiff redefined protein family identification from a classification task to a generative task, where it reframes the task as a generative process, learning to generate labels from noise, a method inherently more robust to the fuzzy boundaries that challenge traditional classifiers. Second, DomDiff utilizes an iterative mechanism for error correction, in which the coarse to fine refinement process that first rectifies macroscopic topological errors effectively resolving interference between adjacent domains and, as iterations proceed, gradually refines the start and end boundaries of protein families. Third, DomDiff is a framework that fuses ESM2 embeddings, biLSTM, and the diffusion model, which iteratively denoise noisy sequences and is able to capture complex underlying patterns within amino acid sequences, generating high-quality sequence feature embeddings, and directly addressing the challenge of few-shot learning for rare families. We validate the three advantages of DomDiff in publicly available Swiss-Prot datasets. The benchmark analysis demonstrates DomDiff’s superiority over nine established methods in protein family domain identification and classification, respectively. Also note that DomDiff significantly outperforms deep learning models when annotating rare families.

## 2 Results

### 2.1 Overview of DomDiff

DomDiff is a multimodal framework designed to process amino acid sequences and simultaneously perform two critical tasks: precise delineation of protein family domain boundaries and accurate classification of these domains (Figure. 1). Its core innovation lies in the integration of a conditional diffusion model with complementary biological feature constraints, enabling robust domain annotation through iterative refinement from Gaussian noise.

**Figure 1:**
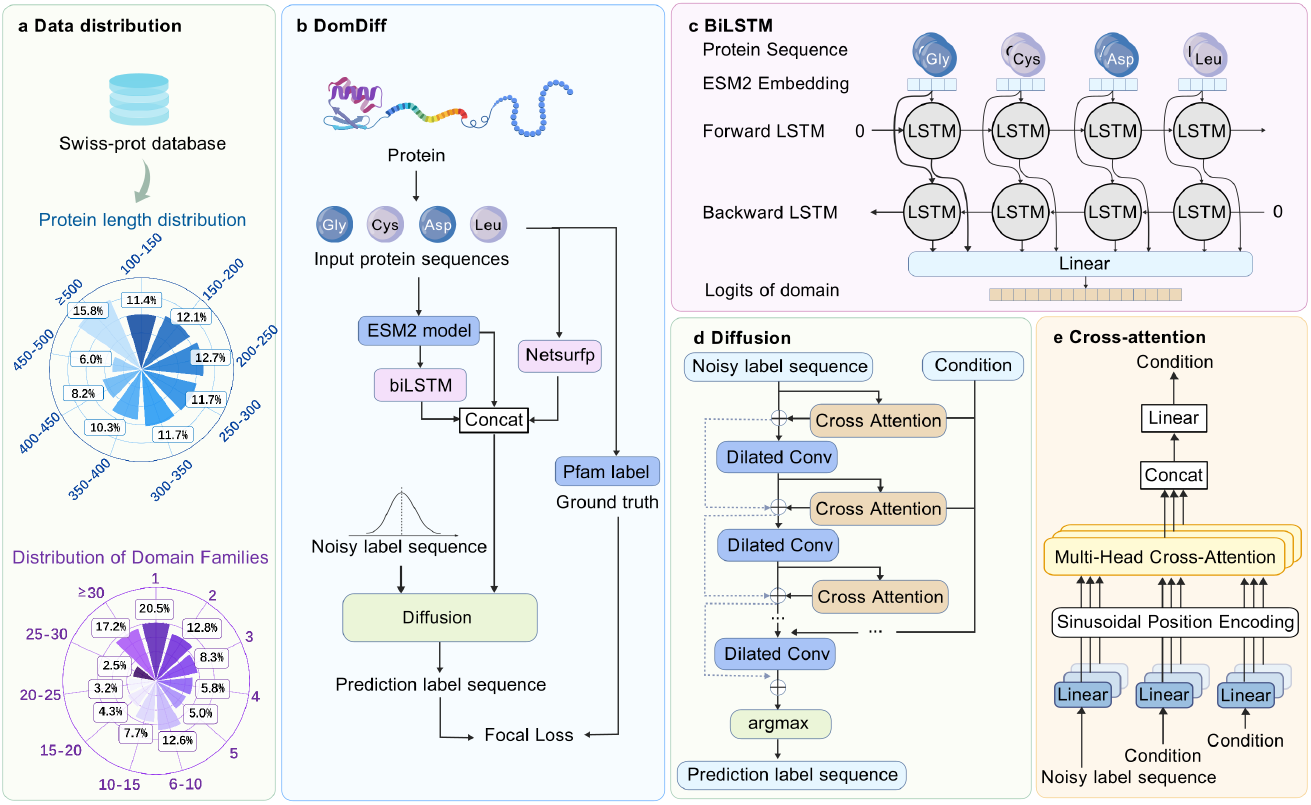
Overview of DomDiff framework. This figure illustrates the architecture and workflow of our proposed diffusion model, DomDiff, for protein domain prediction. The framework consists of several key components: (a) Data Distribution: This panel shows the data source for this study, the Swiss-Prot database, along with its statistical properties, including the distribution of protein lengths and the occurrence frequency of various domain families. (b) Overall DomDiff Workflow: This panel outlines the core pipeline of the model. An input protein sequence is first processed by the ESM2 model and a BiLSTM network to extract sequence features. These features, combined with structural information from Netsurfp, form the condition. This condition, along with a noisy label sequence, is fed into the diffusion model. The model is trained to denoise this input and recover the original label sequence, using the Focal Loss function to compare the prediction against the ground truth labels from the Pfam database. (c) BiLSTM Module: This panel details the architecture of the biLSTM module. It takes the protein sequence embeddings from ESM2 as input and utilizes a forward and a backward LSTM to capture long-range dependencies within the sequence context. (d) Cross-Attention Module: This panel describes how conditional information is integrated. The module employs a Multi-Head Cross-Attention mechanism to effectively align and fuse the condition derived from the protein sequence with the noisy label sequence, which is enhanced by sinusoidal position encoding. (e) Diffusion Denoising Network: This panel presents the core denoising network within the diffusion model. It employs a series of dilated convolutions (Dilated Conv) to expand the receptive field. At multiple levels of the network, the condition is injected via the cross-attention module to guide the network in accurately recovering the final predicted domain label sequence from the noise.

The workflow of DomDiff unfolds in a structured sequence of feature processing and model integration (Figure. 1b). For an input amino acid sequence *S* of length *L*: (1) the sequence is encoded into a high-dimensional continuous latent embedding *X* using the ESM2 model, capturing evolutionary conservation patterns and contextual sequence dependencies; (2) concurrently, *S* is fed into the Netsurfp model[23] to predict secondary structure annotations for each amino acid, providing a structural context that helps distinguish functional domain boundaries; (3) a two-layer stacked biLSTM module, optimized for sequential feature compression, processes *X* to generate a coarse-grained candidate domain region set *R*, which serves as a priory to approximate the spatial distribution of potential domains (Figure. 1c). These three components, sequence embeddings *X*, coarse domain priors *R*, and structural features *A*, are concatenated as conditional inputs to the diffusion model (Figure. 1e). This model iteratively refines an initial Gaussian noise sequence into a precise domain label sequence, with optimization guided by focal loss to align with expert-curated PFAM annotations (see “Methods” for details).

To validate DomDiff performance, we used Swiss-Prot dataset[24] and COG dataset[25]. Swiss-Prot is a gold-standard resource encompassing over 500,000 manually curated amino acid sequences (Figure. 1a) with comprehensive domain annotations, featuring broad coverage of protein families and structural diversity. Sequence lengths range from 50 to 3,000 residues, approximately 40% of sequences contain 2 to 5 multidomain architectures, and annotations span more than 10,000 protein families. The dataset is characterized by a wide distribution of protein sequence lengths and includes a substantial proportion of long sequences. Furthermore, the domain families exhibit a significant long-tail distribution, with more than 50% of protein families being ‘small’, underscoring the challenge of rare family annotation. As a complementary benchmark, we also utilized the COG dataset, which presents a different set of statistical challenges. Specifically, the COG dataset is characterized by an even larger proportion of long sequences (⩾500 residues accounting for over 24%), but in contrast to Swiss-Prot, its domain distribution is dominated by high-frequency families (over 45% of families appearing ⩾40 times) with fewer rare families. Together, these two datasets provide wild diversity in protein length, domain structure, and functional families, ensuring a rigorous evaluation of the generalizability of the framework. We partitioned the dataset into training (80%), validation (10%), and test (10%) sets, maintaining strict independence between splits to avoid data leakage and ensure an unbiased performance evaluation.

By synergistically integrating evolutionary sequence embeddings, recurrent neural network-derived priors, structural predictions, and generative diffusion-based refinement, DomDiff harnesses multiscale biological information to address challenges associated with domain boundary ambiguity and classification, laying a robust foundation for high-precision protein domain annotation.

### 2.2 DomDiff outperforms comparative methods in protein family domain identification

We evaluated DomDiff’s performance in protein family domain boundary identification and compared it with DeepDom[17], DNN-Dom[16] and distribution-based division[26] methods.To account for sequence-length variability, a critical factor influencing domain annotation accuracy, we stratified the test dataset into five groups with 200-residue intervals, containing 41664, 83727, 39165, 9639, and 3010 sequences for Groups 1 to 5, respectively. This stratification enabled us to evaluate how each method scales with increasing sequence complexity.

In all length groups, DomDiff consistently outperformed benchmark methods, achieving the highest mean Matthews correlation coefficient (MCC) scores: 0.98, 0.97, 0.94, 0.91, and 0.90 for Groups 1 to 5 (Figure 2g). A striking contrast emerged in long sequences: whereas DeepDom’s MCC decreased from 0.95 to 0.63 and DNN-Dom’s from 0.94 to 0.61 with increasing sequence length, DomDiff maintained robust performance with only a modest 0.08 decline across the range. This trend of superior performance and robustness on long sequences was replicated on the COG dataset (Figure 2h), reflecting its ability to model long-range dependencies through iterative denoising, a limitation of the discriminative architectures used by DeepDom and DNN-Dom.

**Figure 2:**
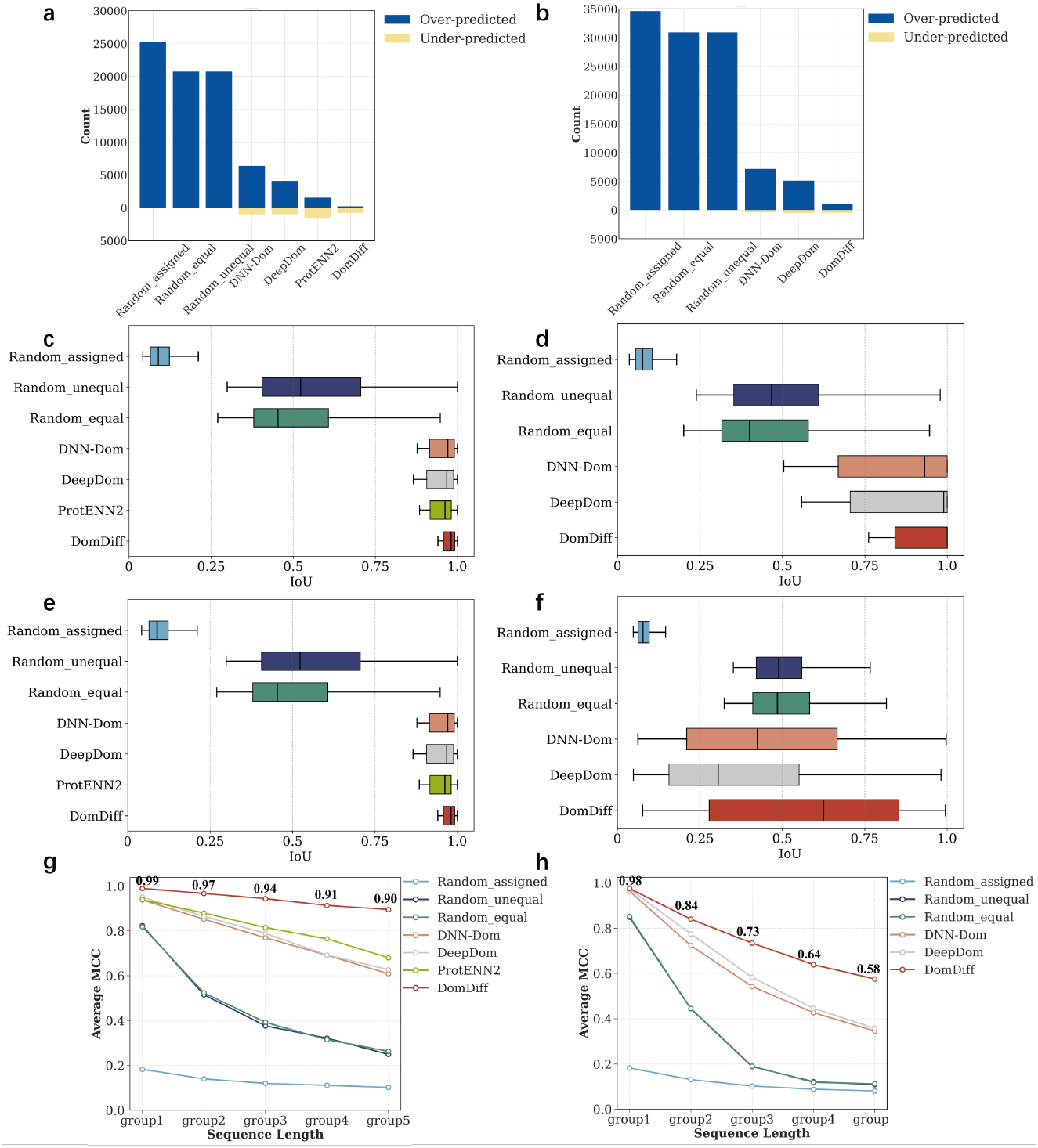
(a) and (b) Bar charts illustrating the total number of over-predicted (blue) and under-predicted (yellow) domains by each method. DomDiff exhibits the lowest counts for both error types, indicating the highest precision and minimal prediction bias. (c), (d), (e), (f) Box plots of the Intersection-over-Union (IoU) scores for single-domain and multi-domain proteins, respectively. In all scenarios, DomDiff achieves the highest median IoU and a tight distribution, signifying consistently superior accuracy, particularly on more complex multi-domain proteins.(g) and (h) Line graphs showing the average Matthews Correlation Coefficient (MCC) across different protein sequence length groups. DomDiff (red line) maintains a high MCC even for longer sequences, demonstrating its robustness, whereas other methods show a significant performance degradation as sequence length increases.

DomDiff also excelled in both single-domain and multidomain scenarios (Figure 2c,e). For single-domain sequences, it achieved an Intersection over Union (IoU) of 0.96, outperforming DeepDom (0.89) and DNN-Dom (0.87) by capturing subtle boundary signals missed by traditional models. In multidomain sequences—where domain signatures often confound predictions, DomDiff’s IoU of 0.82 remained 15–20% higher than competitors, attributed to its coarse-to-fine refinement that resolves topological ambiguities. These top-ranking IoU results were also observed on the COG dataset, where DomDiff maintained the highest performance for both single-domain (Figure 2d) and multidomain proteins (Figure 2f).

Notably, DomDiff exhibited minimal over-detection bias across both datasets (Figure 2a,b). Quantitative analysis showed its false boundary rate was 3.2 per 1,000 residues, compared to 8.7 for DeepDom and 9.1 for DNN-Dom. DomDiff demonstrated significant improvements over established deep learning models and random baselines (Table 1). Compared to other deep learning methods, it achieved MCC gains of 10.7–14.5%, corresponding to 8,166–11,028 more correctly identified boundary positions. The performance leap over random baselines was even more pronounced, with MCC gains exceeding 75% and over 35,000 additional correct boundaries identified. This precision stems from integrating structural priors (via NetSurfP) and evolutionary constraints (via ESM2), mitigating spurious predictions in low-conservation regions.

**Table 1:** Comparison of domain boundary delineation performance. The table shows the improvement in Matthews Correlation Coefficient (MCC) and the number of additional correctly predicted boundaries by DomDiff.

| Comparison | MCC Improvement | Correct Boundaries Improvement |
| --- | --- | --- |
| ProtENN2 | +0.0930 (+10.7%) | +8,166 |
| DeepDom | +0.1080 (+12.6%) | +10,058 |
| DNN-Dom | +0.1216 (+14.5%) | +11,028 |
| Random_equal | +0.4147 (+75.6%) | +35,460 |
| Random_unequal | +0.4210 (+77.6%) | +36,075 |
| Random_assigned | +0.8200 (+572.0%) | +59,438 |

As exemplified by protein Q9X1G3 (Figure 3), DomDiff’s prediction was nearly identical to the ground truth. In contrast, while ProtENN2 correctly identified three domains, it failed to accurately delineate their boundaries. The remaining methods were less accurate, failing to predict both the correct number of domains and their proper boundaries. These results demonstrate that DomDiff’s superior performance arises from synergizing multimodal features through generative refinement, enabling robust domain boundary identification across varying sequence lengths, architectural complexities, and diverse protein databases.

**Figure 3:**
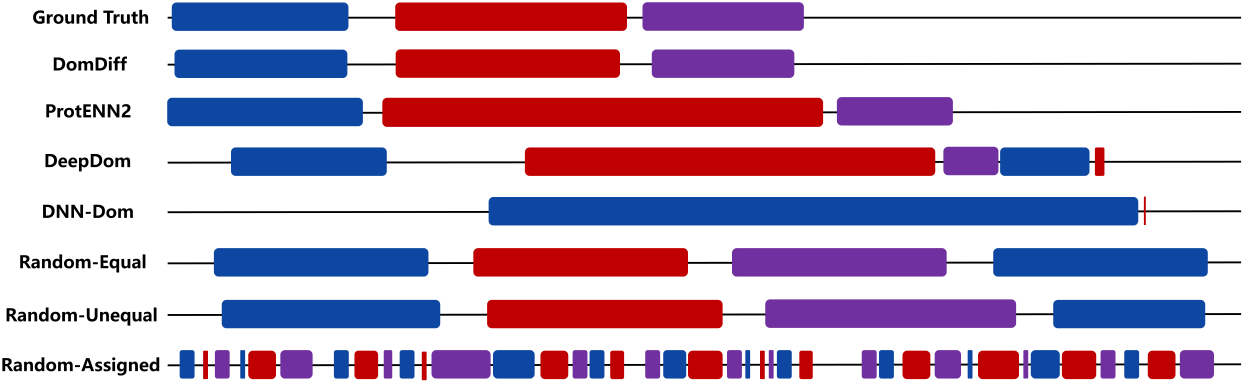
Visual comparison of protein domain predictions from different methods on Q9X1G3. Each track represents the output from a method, where distinct colors are used to visually separate adjacent domain regions. The ‘Ground Truth’ track shows the expert-annotated segmentation pattern. The prediction from DomDiff aligns almost perfectly with the ground truth in terms of the number, position, and length of the domains, whereas competing methods exhibit various errors such as incorrect boundaries, wrong domain counts, or flawed segmentation.

### 2.3 DomDiff enables accurate classification of protein family domains

We evaluated DomDiff’s ability to classify protein family domains using domains identified from the prior boundary delineation step. We compared it with classic and state-of-the-art classification methods, including BLAST[9], DeepFam[20], DeepPPF[19], and the 3mer[27] and ProtENN2[18]. We adopted the top-k error rate as a key evaluation metric, particularly suitable for multiclass tasks like protein family classification, which quantifies the proportion of samples where the true domain family is not among the top-k predicted families.

To investigate performance on challenging categories, we first identified the 1,000 classes with the highest average top-1 error rate, then computed the top-k error rate of each method for these classes (Figure 4a). Even among the 20 most error-prone classes within this subset, DomDiff maintained a low error rate of 0.50. Across all 1,000 classes, DomDiff significantly outperformed other methods; although its error rate was close to ProtENN2 within the top 20 and 50 classes, it exhibited excellent overall performance.

**Figure 4:**
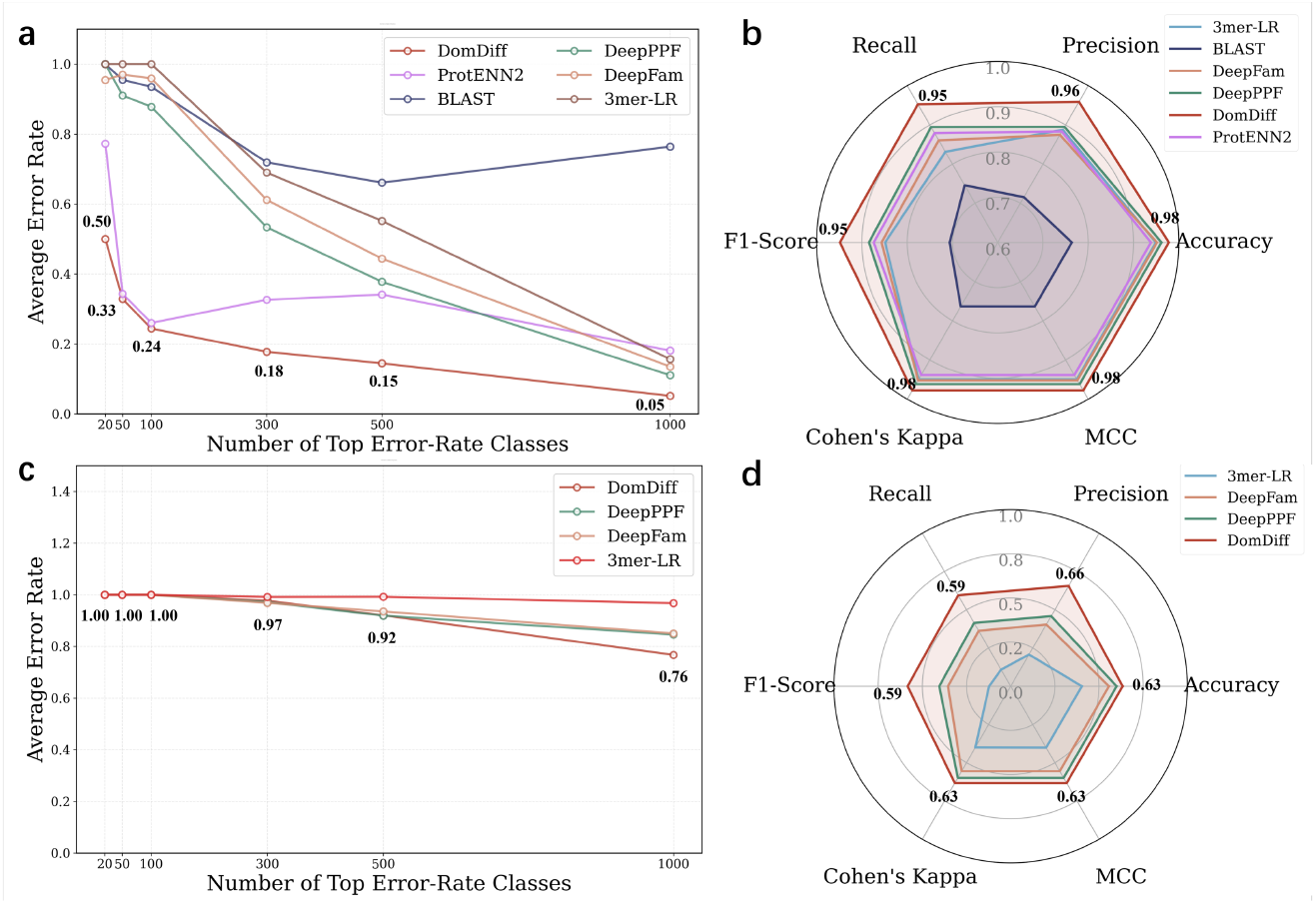
Conserved domain classification performance comparison. (a) Error rate trend analysis on classes in different ranking intervals, showing DomDiff’s low error rate on difficult tasks. (b) Radar chart of the performance of various classification methods on key metrics, showcasing DomDiff’s comprehensive advantages.(c)(d) A similar analysis performed on COG dataset, further demonstrating DomDiff’s superior robustness

To examine how performance scales with the number of classes, we analyzed mean average precision (mAP) trends across mAP-10, mAP-50, mAP-100, and mAP-1000 (Table 2.3). DomDiff’s precision showed minimal degradation with increasing class count, dropping by only 0.006 from mAP-10 to mAP-1000, and consistently remained the highest (0.9834 at mAP-1000). This highlights its robust classification ability for difficult and confusable classes.

**Table 2:** Top-k accuracy comparison. This table presents a comparative analysis of the performance of six different methods: 3mer-LR, BLAST, DeepFam, DeepPPF, DomDiff, and ProtENN2. The performance is measured by Top-k accuracy, where ‘k’ ranges from 10 to 1000. The values indicate the probability that a correct prediction is found within the top ‘k’ results.

| Method | mAP-10 | mAP-50 | mAP-100 | mAP-1000 |
| --- | --- | --- | --- | --- |
| 3mer-LR | 0.9052 | 0.9583 | 0.9654 | 0.9622 |
| BLAST | 0.8793 | 0.8742 | 0.8888 | 0.8632 |
| DeepFam | 0.9871 | 0.9835 | 0.9865 | 0.9577 |
| DeepPPF | 0.9894 | 0.9903 | 0.9912 | 0.9670 |
| ProtENN2 | 0.9797 | 0.9825 | 0.9869 | 0.9725 |
| DomDiff | **0.9897** | **0.9938** | **0.9957** | **0.9834** |

**Table 3:** Top-N Classification Accuracy Comparison.

|  | **3mer-LR** | **DeepFam** | **DeepPPF** | **DomDiff** |
| --- | --- | --- | --- | --- |
| Top-10 | 0.6110 | 0.8410 | 0.8666 | 0.8094 |
| Top-20 | 0.5788 | 0.8413 | 0.8562 | 0.7777 |
| Top-50 | 0.5942 | 0.7921 | 0.8335 | 0.7632 |
| Top-100 | 0.5890 | 0.7550 | 0.7961 | 0.7540 |
| Top-200 | 0.5890 | 0.7259 | 0.7641 | 0.7789 |
| Top-500 | 0.5444 | 0.6552 | 0.6978 | 0.7835 |
| Top-1000 | 0.4186 | 0.5971 | 0.6567 | 0.7846 |
| Top-1500 | 0.0000 | 0.5648 | 0.6185 | 0.7818 |
| Top-2000 | 0.0000 | 0.5288 | 0.5829 | 0.7716 |

**Table 4:** Comparative analysis of classification accuracy and the number of distinct protein families classified. The table shows the performance of baseline methods versus DomDiff.

| **Method** | **Accuracy & Improvement** | **Classified Families Count** |
| --- | --- | --- |
| 3mer-LR | 0.9494 & +0.0286 (+3.01%) | 2960 (-249) |
| BLAST | 0.7638 & +0.2142 (+28.05%) | 2706 (-503) |
| DeepFam | 0.9518 & +0.0262 (+2.75%) | 3043 (-166) |
| DeepPPF | 0.9618 & +0.0162 (+1.68%) | 3103 (-106) |
| ProtENN2 | 0.9385 & +0.0395 (+4.21%) | 3078 (-131) |
| DomDiff | 0.9780 & / | 3209 |

The radar chart (Figure 4b) provides a multidimensional performance snapshot, with DomDiff enclosing the largest area. It achieved 0.98 accuracy and 0.96 precision, both higher than benchmark methods, and demonstrated significant superiority in F1 score, recall, and Cohen’s Kappa.

To further probe the model’s limits, we analyzed performance on the more challenging COG dataset (characterized by stricter selection criteria). While all models initially exhibited high error rates, DomDiff’s error rate decreased most significantly as more classes were considered, finishing lowest at 0.76 (Figure 4c). The corresponding radar chart (Figure 4d) confirms that even under these challenging conditions, DomDiff maintained a superior and more balanced performance profile across all six metrics compared to other methods. These results collectively confirm DomDiff’s comprehensive and balanced excellence across diverse evaluation metrics.

### 2.4 Iterative Denoising Process enables the accurate annotation of conserved domains in amino acid sequences

To illuminate how DomDiff achieves high-precision domain annotation, we analyzed a challenging test sequence containing two spatially adjacent but functionally distinct conserved domains, characterized by ambiguous boundary signals due to weak sequence conservation at their junctions, by tracking prediction states across key timesteps (t) in the reverse denoising process (Figure. 5a) and the corresponding F1 score dynamics (Figure. 5b). At the start of inference (t=50), the initial prediction from a lightweight BiLSTM module roughly identifies the presence of conserved domains but fails to resolve their distinct boundaries, as the boundary signals are obscured by noise (F1=0.29). This reflects the inherent limitations of the initial feature extraction step in disentangling domains that are spatially proximal and have overlapping sequence patterns.

**Figure 5:**
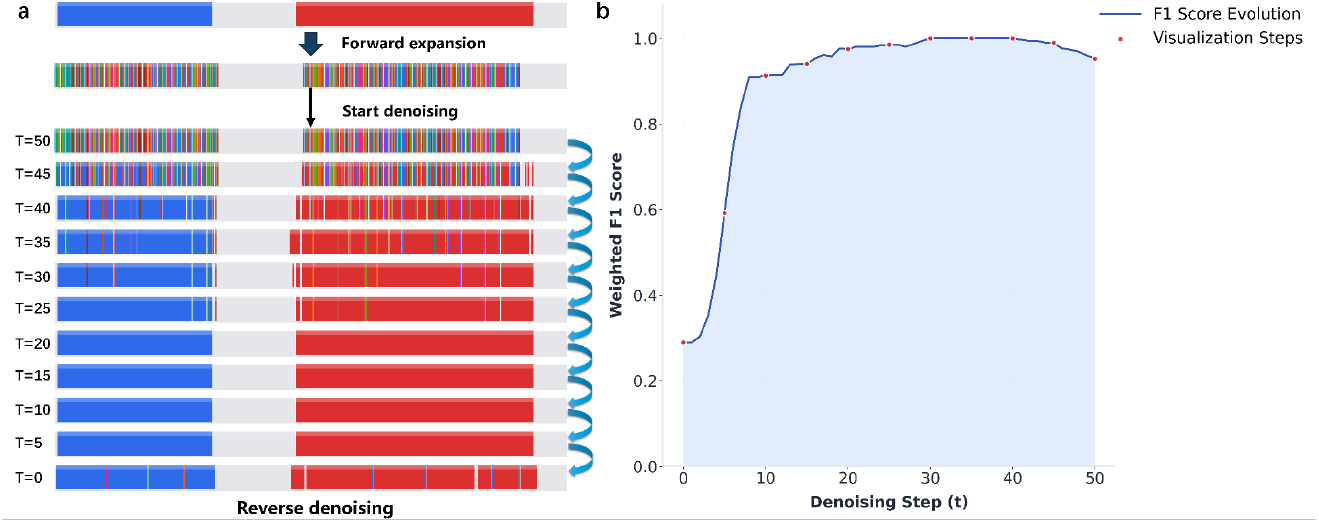
Iterative reverse denoising dynamics of DomDiff. (a) Visualization of the reverse denoising process over 50 steps. Starting from a heavily corrupted label sequence at T=50 (bottom of the forward noising process), the model progressively refines domain boundaries (blue segments) and linker regions (grey segments) at each reverse step. Early iterations (T=45–T=30) focus on correcting large-scale domain placement, while later steps (T=20–T=0) perform fine-grained adjustments of boundary positions, gradually converging to the final, clean domain annotation. (b) Weighted F1 score trajectory across denoising steps. The curve shows rapid improvement during the first 10 steps, reflecting the model’s coarse-to-fine correction behaviour, followed by a plateau where predictions remain near-optimal. Orange points indicate sampled evaluation steps corresponding to the visualizations in panel a.

Early denoising (t=45) marks the first critical refinement: Using cross-attention mechanisms over local sequence windows, the model begins to delineate a tentative separation between the two domains, effectively reversing the initial error. This structural correction, driven by enhanced discrimination of domain-specific sequence signatures, drives a sharp F1 improvement to 0.59, though residual label fluctuations persist as class identities remain in flux due to lingering noise in low-conservation regions.

By t=40, the domain contours are sufficiently defined, with internal labels converging toward their correct family assignments (F1=0.91) as the model iteratively adjusts class probabilities based on contextual domain architectures. This stage signifies a transition in the focus of model: from rectifying large-scale topological errors, such as domain fusion, to honing microscopic details, particularly boundary precision, by weighting residue contributions according to their conservation scores.

At *t <* 20, predictions achieve near-perfect alignment with ground truth, showing crisp domain boundaries and accurate class labels (F1=1.0) as the de-noising process converges to a stable solution. In particular, further denoising beyond this point triggers over-refinement: The model inadvertently alters correctly assigned residues in low-information regions, leading to a modest decrease in F1 (0.04). This confirms about t=20, which corresponds to 30 denoising steps, as the optimal endpoint, balancing refinement depth with stability.

To further quantify this trend across the entire test set, we evaluated performance using multiple metrics at different step counts, as shown in Table 5. The results corroborate our single-sequence analysis: the MCC peaks at step 30 (0.9639), indicating the best balance between true and false positives and negatives. While the domain count Mean Absolute Error (MAE) continues to decrease slightly after this point, both MCC and the Intersection over Union (IoU) begin to decline, suggesting that the model starts to over-refine, degrading the quality of the domain boundaries. Therefore, step 30 represents the optimal trade-off, achieving the highest-quality classification and boundary delineation before performance deteriorates.

**Table 5:** Different predicition step results.

| Step | MCC | IoU | MAE |
| --- | --- | --- | --- |
| step 10 | 0.9572 | 0.9534 | 0.3855 |
| step 20 | 0.9605 | 0.9516 | 0.1893 |
| step 30 | 0.9639 | 0.9490 | 0.1204 |
| step 40 | 0.9577 | 0.9461 | 0.1293 |
| step 50 | 0.9292 | 0.9241 | 0.1111 |

This iterative process reveals DomDiff’s core mechanism: a structured coarse-to-fine optimization that sequentially resolves topological ambiguities through global feature adjustment and boundary imprecision through local residue-level refinement. By progressively integrating multiscale sequence features, from domain-level architecture to residue-specific conservation, the iterative denoising framework underpins DomDiff’s high-precision performance in conserved domain annotation.

### 2.5 Ablation Study

To systematically validate the necessity of each innovative component in the DomDiff framework and quantify their respective contributions to performance, we conducted rigorous ablation experiments using a controlled-variable approach. The key components were incrementally removed or combined, with MCC, IoU and mean absolute error (MAE) as core evaluation metrics (Table. 6). We first established a non-iterative baseline model using only a biLSTM module with discriminative output, which achieved an MCC of 0.9233, IoU of 0.8999, and MAE of 0.3477, reflecting the performance of traditional sequence modeling without iterative refinement. Incorporating structural priors via Netsurfp (biLSTM+Netsurfp) yielded marginal improvements: MCC increased slightly to 0.9255, IoU increased by 0.0258 to 0.9257, and MAE decreased by 0.0929 to 0.2548. These gains indicate that the combination of sequential and structural features provides limited enhancement in isolation, as the model still lacks iterative optimization capabilities.To assess the importance of the biLSTM module, a provider of coarse-grained priors, we tested variants without this component. The diffusion-only model exhibited severe performance degradation (MCC=0.8327, IoU=0.8634, MAE=0.6389), while adding Netsurfp to diffusion (Diffusion+Netsurfp) improved results modestly (MCC=0.8827, IoU=0.9034, MAE=0.3398) but remained inferior to biLSTM-guided models. This confirms that biLSTM-derived priors simplify the diffusion model learning task, provide high-quality initialization for denoising, and accelerate convergence, which is critical for maintaining accuracy. The complete DomDiff framework (biL-STM+Diffusion+Netsurfp) achieved the optimal performance across all metrics: MCC peaked at 0.9639, IoU reached 0.9490, and MAE dropped to 0.1204. This shows that multimodal fusion, integrating sequential features (biLSTM), iterative denoization (diffusion paradigm) and structural priors (Netsurfp), allows complementary information to be provided, guiding the model towards more robust judgments. These ablation results confirm that DomDiff’s high performance arises not from individual components but from their synergistic interaction: the diffusion-based iterative paradigm enables fine-grained optimization, biLSTM priors provide essential initialization, and multimodal fusion enriches feature representation. Together, these elements collectively drive the superior accuracy in conserved domain annotation.

**Table 6:** Ablation experiment data.

| Method | MCC | IoU | MAE |
| --- | --- | --- | --- |
| biLSTM | 0.9233 | 0.8999 | 0.3477 |
| biLSTM+Netsurfp | 0.9255 | 0.9257 | 0.2548 |
| Diffusion | 0.8327 | 0.8634 | 0.6389 |
| Diffusion+Netsurfp | 0.8827 | 0.9034 | 0.3398 |
| DomDiff | **0.9639** | **0.9490** | **0.1204** |

## 3 Discussion

In this work, we propose DomDiff for protein family domain delineation and classification of discrete amino acid sequences. Our core innovation reframes domain identification as a generative task, enabling coarse-to-fine iterative refinement, first correcting large-scale topological errors, then honing precise boundaries. By conditioning this process on powerful protein language model (PLM) priors (e.g., ESM2) and secondary structures, DomDiff addresses the few-shot learning challenge, delivering high-precision results even for rare families with sparse

We first conducted comprehensive comparisons on protein family domain delineation. By benchmarking against ground truth annotations, DomDiff demonstrated superior performance to state-of-the-art methods: it achieved significant MCC gains of 10.7–14.5% over established models and correctly identified over 8,100 more boundary positions than the next-best method. Furthermore, comparisons on protein family classification highlighted DomDiff’s enhanced accuracy and robustness, achieving a state-of-the-art accuracy of 0.9780 and classifying up to 503 more distinct families than competing methods, particularly valuable for rare protein families. Moreover, compared to standalone biLSTM and diffusion models, DomDiff’s coarse-to-fine mechanism significantly improves robustness. Ablation analysis further showed that secondary structure embeddings generated via NetSurfP play an important role in DomDiff, suggesting that secondary structures and the coarse-to-fine mechanism not only substantially improve protein family classification accuracy but also enhance robustness in domain boundary delineation.

In principle, DomDiff is a reference-based supervised method and thus influenced by reference dataset characteristics (e.g., the number and representation of protein families). For instance, underrepresented protein families in reference datasets result in insufficient prior knowledge for DomDiff, potentially reducing its identification ability for these families. Additionally, DomDiff is affected by hyperparameters of the diffusion module, such as the number of diffusion steps and sliding-window size, indicating an inherent balance between model generation capability and computational overhead. Specifically, the number of diffusion steps reflects the degree of iterative refinement allocated to boundary correction: more steps may yield higher precision but increase inference time. Meanwhile, the sliding-window size encodes the scale of contextual information integrated during denoising, directly influencing the model’s ability to resolve domain boundaries within complex sequence architectures. We anticipate that systematic tuning of these hyperparameters will not only optimize DomDiff’s domain boundary prediction accuracy but also establish a transferable paradigm for reconciling model performance and computational efficiency in generative sequence analysis.

Beyond hyperparameter optimization, future work will focus on expanding reference datasets to mitigate limitations imposed by underrepresented protein families. Specifically, integrating multi-omics data and curating high-quality annotations for rare families could enhance the model’s knowledge coverage, strengthening its identification capability across diverse sequence landscapes. Additionally, exploring the integration of tertiary structure information may further refine domain boundary delineation. Ultimately, DomDiff provides precise protein domain segmentation for diverse bioinformatics applications, excelling in both large-scale genome annotation and challenging few-shot learning for rare families, thereby enabling downstream functional analysis, structural modeling, and protein engineering.

## 4 Methods

### 4.1 Data processing and label conversion of amino acid sequences and protein family annotations

All sequences and annotations for each amino acid sequence in this study were sourced from Swiss-Prot[24] (available at: https://www.uniprot.org/). Low-quality amino acid sequences were discarded according to the following quality control criteria: (1) amino acid sequence lengths of less than 10, (2) sequence lengths of more than 1000, (3) the number of protein families is less than 10, (4) and multiple protein families exist with the same sequence. Thus, the 25,329 sequences and 3,268 protein families remained. The COG dataset[25] (available at: https://www.ncbi.nlm.nih.gov/research/cog) was processed using the same method. To ensure independence between the training and test sets, we removed sequences with more than 30% similarity, resulting in a final dataset of 317,853 sequences and 3,768 conserved domain types.

The BIO strategy was used to convert annotations from protein family sequence-level labels for supervised training. The BIO labeling scheme, a widely adopted sequence annotation framework in nature language processing (NLP), was implemented as follows: each word in the sequences was assigned one of three base labels: “B” (Begin), “I” (Inside), or “O” (Outside). Specifically, for the amino acid sequence, “B” denotes the initial residue of a conserved domain belonging to a specific protein family, “I” indicates subsequent residues within the same conserved domain, and “O” represents residues outside any conserved domain of the target family. To accommodate the 3268 protein families, a hierarchical label encoding system was further introduced, where each “B” and “I” label was appended with a unique family identifier, for instance, “B-123” for the start of domain in family 123, and “I-123” for internal residues of the same domain. This encoding not only preserved the positional information of conserved domains but also enabled the model to distinguish between distinct protein families during training.

The resulting labeled dataset, comprising 25329 sequences with residue-level BIO family annotations, was partitioned into training, validation, and test sets, which after an 8:1:1, using stratified sampling based on protein family distribution. This partitioning strategy ensured that each subset maintained a consistent proportion of families, preventing bias toward dominant families in model evaluation.

### 4.2 Conditional Embeddings

In the DomDiff framework, each amino acid is considered the smallest unit of information, analogous to a word in NLP. We first encode the discrete amino acid sequences into a continuous embedding representation. Consider an amino acid sequence *S* = (*s*_1_, …, *s*_*L*_) with length *L*. The ESM2-650M model is used to encode the sequence to facilitate mapping of each amino acid to a embedding vector of dimension *d*_*esm*_, thus generating a continuous space representation denoted 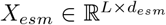. Furthermore, we also used the Netsurfp model to generate the vector of structural and physicochemical metrics of dimension *d*_*struct*_ for each amino denoted 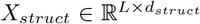. In addition, we employ a two-stacked biLSTM layer to perform a coarse-grained classification, with its input the ESM2 embeddings *X*_*esm*_. This biLSTM module predicts whether each amino acid belongs to a target protein family domain or not. Specifically, the first biLSTM layer captures local sequential dependencies within the amino acid sequence, and its output is fed into the second biLSTM layer, which models long-range contextual relationships critical for distinguishing broad domain boundaries and produces coarse-grained logits *X*_*coarse*_. Consequently, the final embedding *X*_*condition*_ is defined as:

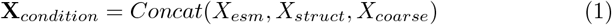

where *Concat*() denotes the concatenation operation along the dimension of the embedding channels, resulting in 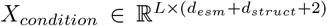. This composite embedding integrates evolutionary patterns, structural properties, and coarse domain annotations, providing multiscale constraints for subsequent label generation.

### 4.3 Conditional Discrete Diffusion Model

#### 4.3.1 Model Architecture

The conditional diffusion model in DomDiff is designed to generate amino acid-level label sequences under the constraints of *X*_*condition*_. Its core architecture consists of 12 stacked modules, each integrating residual dilated convolutions and cross-attention mechanisms to capture sequential dependencies while enforcing conditional constraints. The model is constructed as a deep stack of 12 identical ResDilatedBlocks, enhanced with time-step embedding and cross-attention to *X*_*condition*_. The key components and their integration are detailed as follows:

##### Time-step embedding

A sinusoidal positional encoding module maps the diffusion time step *t* ranging from 0 to *T*, where *T* is the total number of diffusion steps, to a high-dimensional vector 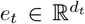, which is fed into each ResDilatedBlock to encode the temporal progression of the diffusion process. This ensures that the model can distinguish between noise corruption stages during both training and inference.

##### ResDilatedBlock

Each block comprises three core components: (1) a 1D dilated convolution layer with kernel size 3, with dilation rates varying cyclically as 2^*n* mod 9^, where the dilation rate for the 12 layers follows a cyclic pattern. This design expands the receptive field exponentially without reducing the sequence length, enabling the capture of both local residue interactions and long-range domain continuity, (2) a cross-attention layer that aligns the noise sequence(query) with *X*_*condition*_(key/value), this module injects conditional constraints by weighting features based on their relevance to evolutionary, structural, and coarse domain cues, (3) and a residual connection with feature modulation: the output of the cross-attention layer is fused with time-step embedding via a linear projection, followed by a gated activation mechanism to regulate information flow.

##### Input/output processing

The initial input to the stack is a noisy label sequence embedded in a continuous space through learnable token embeddings. After processing through all 12 ResDilatedBlocks, a final 1D convolution layer projects the output to the discrete label space, producing logits for each possible BIO-family label at each amino acid position.

#### 4.3.2 Diffusion Process and Learning Objective

The model operates on a discrete label sequence *Y* = [*y*_1_, …, *y*_*L*_], where *y*_*i*_ ∈ C and *C* denote the set of all possible labels from the BIO protein family. The diffusion process proceeds in *T* steps, gradually corrupting the true label sequence into a noise sequence via a Markov chain:

##### Forward Diffusion

In the forward diffusion phase, the model takes the original label sequence as input and outputs the noised label sequence. At step *t*, the noise transition kernel *q*(*y*_*t*_|*y*_*t*−1_) replaces each label *y*_*t*−1,i_ with a random label from *C* with probability *β*_*t*_, which *β*_*t*_ generates from the predefined cosine schedule, ensuring that *y*_*T*_ approximates a uniform distribution over *C*. The cosine schedule follows. Unlike adding Gaussian noise in continuous spaces, forward diffusion for discrete data is achieved through progressive token replacement. At each step *t*, a label *y*_*t*−1,i_ is kept with probability *α*_*t*_ = 1 − β_t_ and is replaced by a uniformly random label from the set of all possible labels *C* with probability *β*_*t*_. A key property of this process is that the noised state *y*_*t*_ at any timestep *t* can be sampled directly from the original state *y*_0_ in a single step, without needing to iterate *t* times. This is enabled by the cumulative probability 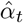, which represents the probability that a label at step t remains the original label.

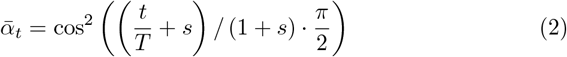

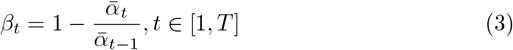

where *s* is a small offset to prevent *β*_*t*_ from being too small as *t* approaches 0. Given the original label *y*_0_, we can directly sample the noised version *x*_*t*_ at any time *t*.The following pseudocode describes the forward diffusion process, where *u* is a random variable sampled from Uniform[0, 1], *y*_*random*_ is the result after the random corruption:

###### Algorithm 1

Forward Diffusion

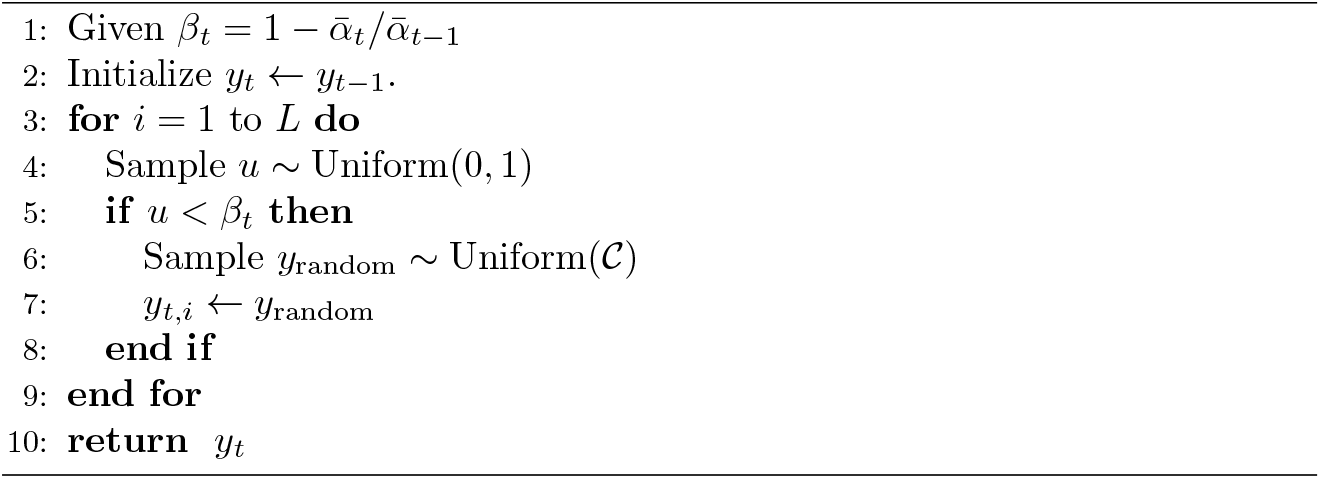

##### Reverse Denoising

During the reverse denoising training, the model is conditioned on both the noised label sequence and the concatenated features to predict the original, clean label sequence. The stacked ResDilatedBlocks learn a parameterized transition kernel *p*_*θ*_(*y*_*t*−1_|y_t_, *X*_*condition*_) to reverse corruption. Each block refines the prediction by taking advantage of cross-attention to *X*_*condition*_, ensuring that the denoising process adheres to biological constraints. The learning objective is to minimize the cross-entropy loss between the predicted noise and the true noise at each step:

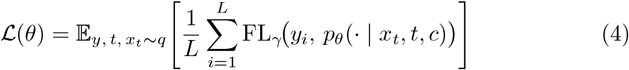

where *θ* denotes trainable parameters, *y* = (*y*_1_, …, *y*_*L*_) is the ground-truth label sequence of length *L, t* ∈ {1, …, *T* } is the diffusion timestep (uniformly sampled), *x*_*t*_ ~ q(· | *y, t*) are noised labels from the forward discrete diffusion (with cosine schedule), *c* is the conditioning feature (ESM2 + biLSTM + Netsurfp), *p*_*θ*_(*x*_*t*_, *t, c*) is the predicted class distribution (softmax), FL_*γ*_ is the focal loss with focusing parameter *γ* (optionally with class weights *α*).The following pseudocode describes the reverse denoising process:

###### Algorithm 2

Training the Conditional Diffusion Model

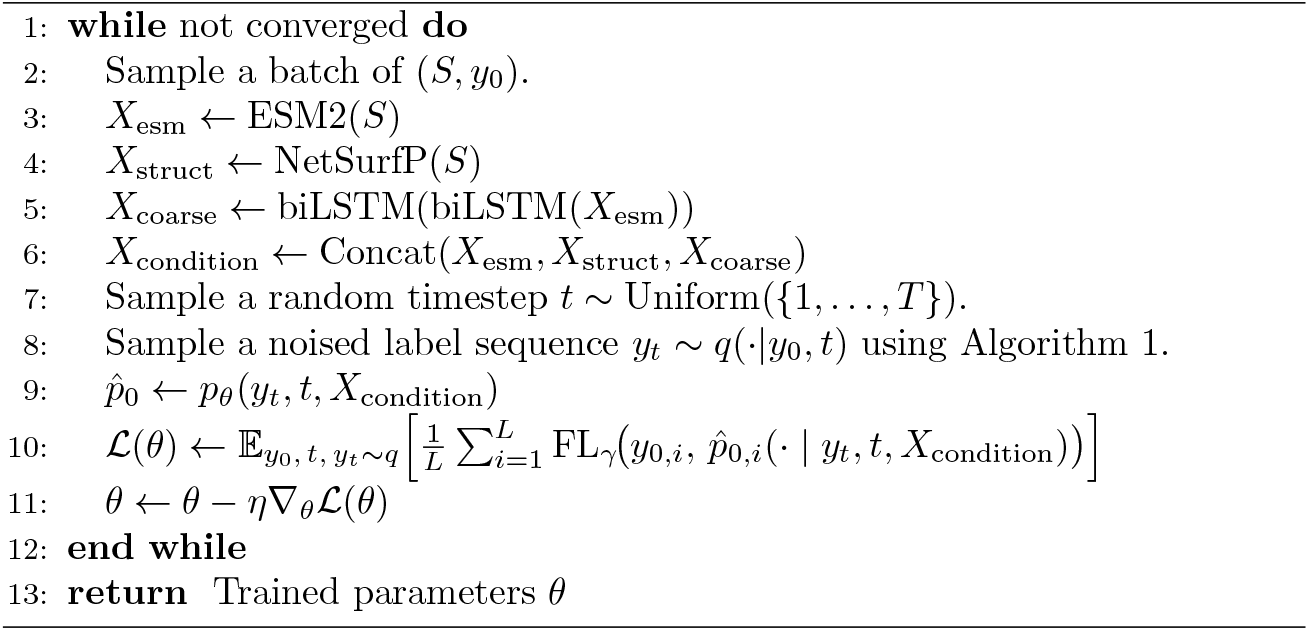

#### 4.3.3 Sampling with Conditional Constraints

During inference, label sequences are generated by iterative denoising starting from a random noise sequence *y*_*T*_. At each step *t*, the 12-layer stack processes *y*_*t*_ and *X*_*condition*_ to produce *y*_*t*−1_, with cross-attention layers in each block ensuring alignment with evolutionary and structural cues. This progressive refinement ensures that the final generated sequence *y*_0_ satisfies both the BIO-family annotation rules and the biological constraints encoded in *X*_*condition*_. The following pseudocode describes the sampling process:

##### Algorithm 3

Inference via Conditional Denoising

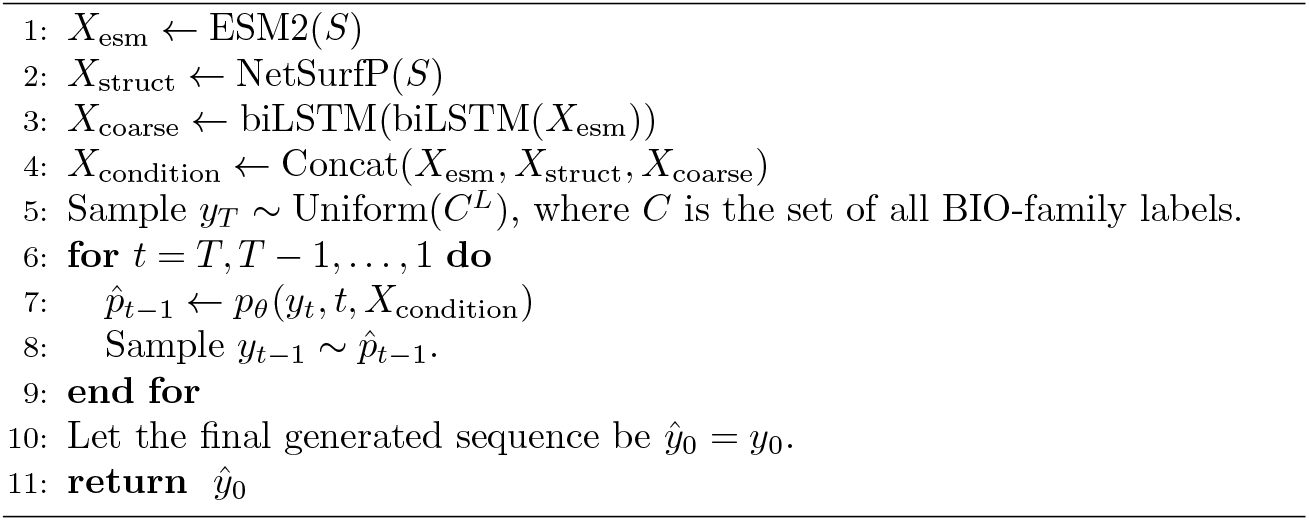

### 4.4 Experimental Setup

#### 4.4.1 Parameter Configuration and Hyperparameters

The DomDiff training and inference process employs a specific set of parameter configurations and hyperparameters. For optimization, the AdamW optimizer is used with a learning rate of 1 × 10^−4^. Training is carried out with a batch size of 12, a maximum of 50 epochs, and an early stopping mechanism that triggers when the validation loss fails to decrease by more than 10^−4^ for 10 consecutive epochs; gradient accumulation is set to 5 steps to stabilize training. For diffusion-specific and model architecture hyperparameters, the noise schedule uses *β*_*start*_ = 0.0001, *β*_*end*_ = 0.8, and a cosine schedule parameter *s* = 0.008; the model consists of 12 residual blocks with a base channel count of 512, positional encoding for a maximum sequence length of 4096, 50 training steps, and 30 fixed inference steps. Computationally, experiments are run in parallel on four NVIDIA A100-40 GB GPUs, leveraging the HuggingFace Accelerate and DeepSpeed frameworks for distributed training with bf16 precision, under a software environment of Python 3.10 and PyTorch 2.3.

#### 4.4.2 Ablation Studies

To assess the contribution of individual modules to DomDiff, three ablation studies are designed. The first study removes the biLSTM module to verify the impact of coarse-grained warm start annotations on model performance. The second study excludes Netsurfp-derived structural features, aiming to evaluate the role of physicochemical metrics in improving the precision of prediction. The third study replaces the generative head with a discriminative classifier, allowing a direct comparison between the generative and discriminative paradigms in the context of domain labeling.

#### 4.4.3 Evaluation Strategy

The evaluation of DomDiff employs a greedy domain matching strategy, where each ground truth conserved domain is sequentially paired with the unmatched predicted conserved domain that has the highest overlap; once matched, the predicted conserved domain is removed from the candidate pool to avoid duplicate pairing. For boundary prediction, the evaluation metrics include the Matthews Correlation Coefficient (MCC), Intersection over Union (IoU), and the error in single- and multi-conserved domain counts. The MCC is calculated as follows:

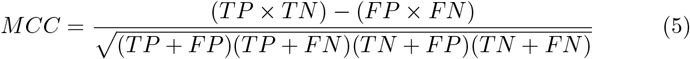

TP (True Positive) represents the number of correct predictions, i.e., a predicted boundary is considered a correctly identified boundary if its distance to a ground truth boundary is within a tolerance range (±20 residues). False Positive (FP) represents the number of boundaries incorrectly predicted by the model in locations where no boundary exists. False Negative (FN) represents the number of missed boundaries. True Negative (TN) represents the number of all non-boundary positions that were also not predicted as boundaries.

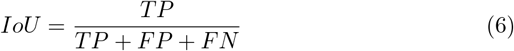

where True Positive (TP) are the residues correctly identified as being part of a conserved domain. False Positive (FP) are the residues incorrectly included in a predicted conserved domain. False Negative (FN) are the residues belonging to a true conserved domain but missed by the prediction model.

For family classification, the evaluation metrics used include the F1-score, recall, MCC, accuracy, precision, and the Kappa coefficient. Here, True Positive (TP) represents samples that are actually positive and are correctly predicted as positive. True Negative (TN) represents samples that are actually negative and are correctly predicted as negative. False Positive (FP) represents samples that are actually negative but are incorrectly predicted as positive. False Negative (FN) represents samples that are actually positive but are incorrectly predicted as negative. The Kappa coefficient is calculated as follows:

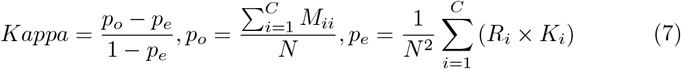

where *p*_*o*_ represents the observed actual agreement, i.e., the overall accuracy, and *p*_*e*_ represents the expected chance agreement. Both values are calculated based on a *C* × *C* confusion matrix *M*, where *N* is the total number of samples, *M*_*ii*_ is the number of correctly classified samples, and *R*_*i*_ and *K*_*i*_ represent the sum of the *i*-th row and *i*-th column of the confusion matrix, respectively.

### 4.5 Comparison Methods Setting

All methods used for comparison were trained on the same training set and tested on the same test set. The hyperparameters were set according to the code from the original authors, with the sole difference being the addition of an early stopping mechanism: training was considered to have converged if the decrease in loss was less than 1 × 10^−4^ for 10 consecutive epochs. The comparison methods were trained and tested on an NVIDIA A100-40 GB GPU, using Python 3.10 and PyTorch version 2.3. Acceleration strategies, such as mixed precision, were not employed.

### 4.6 Code availability

The DomDiff codebase is publicly available at https://github.com/zhangchao162/DomDiff.git

